# Impute.me: an open source, non-profit tool for using data from DTC genetic testing to calculate and interpret polygenic risk scores

**DOI:** 10.1101/861831

**Authors:** Lasse Folkersen, Oliver Pain, Andres Ingasson, Thomas Werge, Cathryn M. Lewis, Jehannine Austin

## Abstract

To date, interpretation of genomic information has focused on single variants conferring disease risk, but most disorders of major public concern have a polygenic architecture. Polygenic risk scores (PRS) give a single measure of disease liability by summarising disease risk across hundreds of thousands of genetic variants. They can be calculated in any genome-wide genotype data-source, using a prediction model based on genome-wide summary statistics from external studies.

As genome-wide association studies increase in power, the predictive ability for disease risk will also increase. While PRS are unlikely ever to be fully diagnostic, they may give valuable medical information for risk stratification, prognosis, or treatment response prediction.

Public engagement is therefore becoming important on the potential use and acceptability of PRS. However, the current public perception of genetics is that it provides ‘Yes/No’ answers about the presence/absence of a condition, or the potential for developing a condition, which in not the case for common, complex disorders with of polygenic architecture.

Meanwhile, unregulated third-party applications are being developed to satisfy consumer demand for information on the impact of lower risk variants on common diseases that are highly polygenic. Often applications report results from single SNPs and disregard effect size, which is highly inappropriate for common, complex disorders where everybody carries risk variants.

Tools are therefore needed to communicate our understanding of genetic predisposition as a continuous trait, where a genetic liability confers risk for disease. Impute.me is one such a tool, whose focus is on education and information on common, complex disorders with polygenetic architecture. Its research-focused open-source website allows users to upload consumer genetics data to obtain PRS, with results reported on a population-level normal distribution. Diseases can only be browsed by ICD10-chapter-location or alphabetically, thus prompting the user to consider genetic risk scores in a medical context of relevance to the individual.

Here we present an overview of the implementation of the impute.me site, along with analysis of typical usage-patterns, which may advance public perception of genomic risk and precision medicine.

## 1 Introduction

In clinical genetics, testing for rare strong-effect causal variants is routinely performed in the healthcare system to confirm a diagnosis, or to evaluate individual risk suspected from anamnestic information [Baig et al 2016], and in such instances the use of genome sequencing is expanding [Byrjalsen et al 2018]. Meanwhile, outside of the healthcare system, direct to consumer (DTC) genetics expands rapidly providing the public with access individual genetic data profiles and to interpretation of common genetic variants derived from genotyping microarrays. This is developing as a sprawling industry of consumer services with widely diverging standards, including third-party genome analysis services. These services typically provide individual results from analysis of single, common SNPs with (at best) weak effects. They are therefore severely mis-aligned with current state-of-the-art, which at least for common, complex disease is to use polygenic risk scores (PRS) to estimate the combined risk of common variation in the genome [Lewis et al 2017].

We believe that the goal of the academic genetics community should extend beyond theory. This means engaging with the public and assisting those that seek information, even when it means helping them to interpret their own genomic data. We therefore developed impute.me as an online web-app for analysis and education in personal genetic analysis. The web-app is illustrated in figure 1. Using any major DTC vendor, a user can download their raw data and then upload it at impute.me. Uploaded files are checked and formatted according to procedures that have been developed to handle most types of microarray-based consumer genetics data, including an imputation step. This data is then further subjected to automated analysis scripts including polygenic risk score calculations. This includes more than 2000 traits, browsable in different interface-types (modules). Each module is designed with the goal of putting findings in as relevant a context as possible, prompting users to see common variant genetics as a support tool rather than a diagnosis finder. The aim is to provide information as broadly as possible to offer a real alternative to the wide-spread practice of reporting on weak single SNP genotypes for any trait, even though that entails generation of some reports that are below any sensible threshold for clinical usability. We hope that having this as an open and accessible resource for everyone will be of help to the debate on what exactly constitutes clinical usability beyond high-risk pathogenic variants.

**Fig 1 –.**
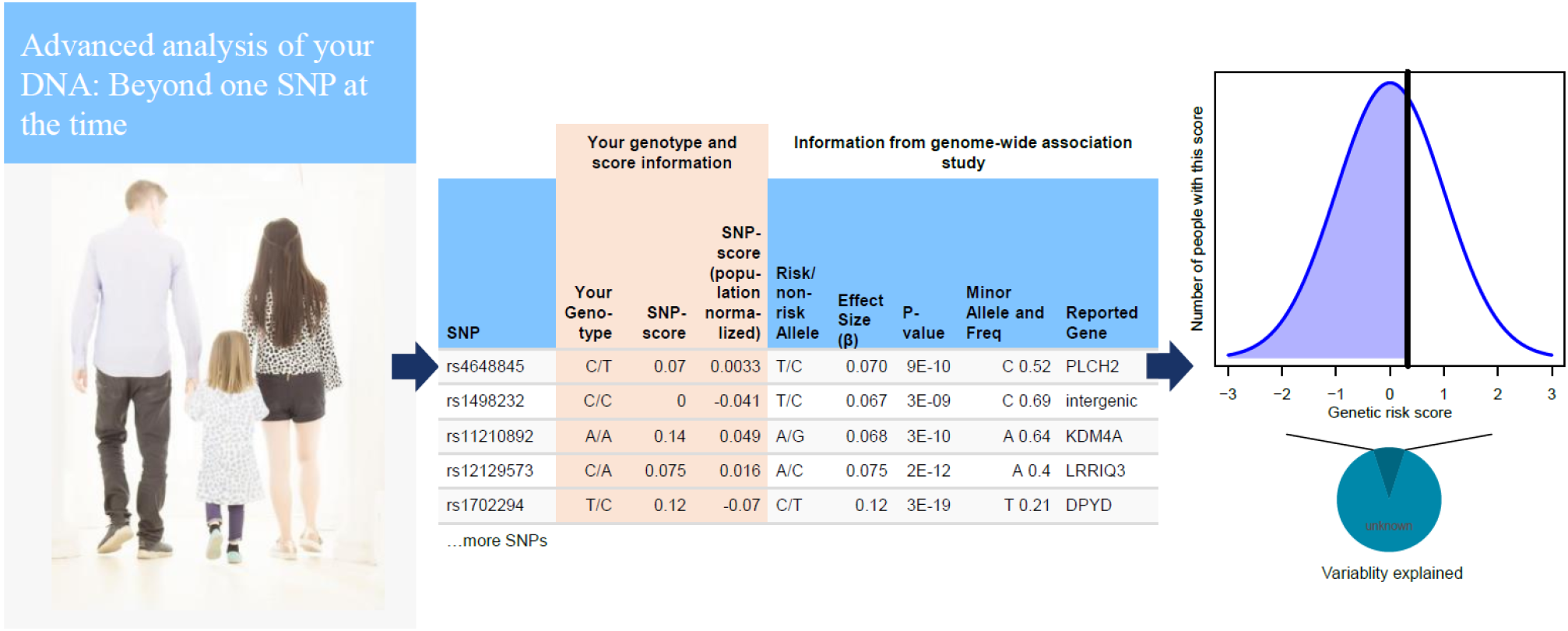
basic pipeline setup from the user point of view. On upload of a genome, data is checked according QC parameters that have been developed to handle most types of microarray-based consumer genetics data. The genome is then imputed using 1000 Genomes as reference (**left**). The imputed data is then further subjected to automated analysis scripts from 15 different modules, most of which are based on polygenic risk score calculations. The calculations include 1859 traits from GWAS studies, 634 traits from the UK Biobank, as well as customized modules for height, and drug response. Most polygenic risk scores use GWAS significant SNPs out of necessity, although 20 major diseases are based on LDpred all-SNP scores (**center**). User can then browse their scores in relation to the population, shown together with a chart displaying how much varability is explained (**right**).

In this paper we will describe the i) development and setup, ii) validation and testing and iii) evaluation of usage, and iv) future directions for impute.me. In the section *Development and Setup* we discuss some of the challenges faced when developing a full personal-genome scoring pipeline. The goal of this section is to motivate and explain the choices made in development. In the second section, *Validation and Testing*, we use public biobank data from individuals that are consented for genetic research to test the effect of the impute.me scores on known disease outcomes. The purpose of this section is to test and validate scores, as well as to investigate consequences of some of the challenges that were raised in the first section. In the third section, *Evaluation of Usage*, we evaluate usage-metrics of impute.me users. The goal of this section is to shed light on behaviour patterns of individuals who use DTC genetics for health questions and offer recommendations that may be of use in other personal-genome scoring pipelines. Finally, in the section *Future Directions we* discuss our views on future directions particularly with respect to improving how genetic findings are presented to people.

## 2 Development and setup

A first challenge in development of personal genomic services is standardization. As the name impute.me implies, all genotype data is processed by imputation of genotype data [Delaneau et al 2013, Howie et al 2009]. This procedure expands the data available into ungenotyped SNPs and increases overlap with public genome-wide association study (GWAS) summary statistics used to estimate risk. It also expands the SNP overlap between microarray types from the major vendors, such as 23andme, Myheritage and Ancestry.com. Further, we have found that imputation helps in avoiding major errors, for example strand-flip issues that arise from the dozens of different data formats. Eliminating such problems from further processing is one important step to minimize mis-interpretation of genome analysis. To ensure high standard of reported results, impute.me requires a fully completed imputation for continued analysis.

The second challenge is calculation of robust PRS estimates that are accurate, irrespectively of the source of the data. This is particularly important to an application utilized by people from around the world leveraging data from dozens of different vendors and data types. Importantly, PRS calculated from GWAS of a population of (e.g.) European ancestry, will perform better for individuals of the same ancestry, and the systematic shift (i.e. bias) in risk scores in individuals from other populations is a problem [Curtis 2018]. Since studies of all disease-traits are not yet available for all non-European populations, the pragmatic solution has been to include a population-specific normalization attempting to minimize the systematic shifts of scores for non-European ancestry users. Further, it is computationally and logistically easier to implement PRS that use only the most (i.e. genome-wide) significant SNPs (often referred to as top-SNPs), but the prediction strength is better when more SNPs are included (all-SNP) which however, is more sensitive to ethnic biases [Lam et al 2019]. The impute.me pipelines calculate PRS for each trait or disease based on all-SNP based PRS calculations if full genome-wide summary statistics are available and processed, and top-SNP based PRS calculations if not.

The third challenge is presentation. For a single rare large-effect variant, such as for the pathogenic variants in the *BRCA* genes conferring very high risk of cancers (odds-ratio >10; Figure 2A, upper left), presentation focuses on absence-vs-presence [Maxwell et al 2016]. However, also low-effect variants, e.g. as in pharmacogenetics, impacting on statin-response, is considered as having potential clinical use [Natarajan et al 2017] (figure 2A, lower right). This difference in effect-magnitude is a major challenge in result presentation, and the line between useful and not depends on context. In the context of a drug-prescription situation or a question of which of two suspected disease risks are the most likely, it may be useful to know such scores. But in the context of an otherwise healthy individual, genetic risks are only relevant if we are very certain of them, they are serious, and preferably actionable (e.g. BRCA variants [Kalia et al 2017]). According to this, that risk-types inherently are not equal, a design choice is that impute.me does not use risk-sorted lists. Currently scores are accessible either through an alphabetically sorted list or in a tree-like setup where genetic scores are reported in a health-context-tree (figure 2B). In this, all scores are included, but scores that are less relevant to healthy individuals (i.e. most of them), are buried deeper into the healthcontext-tree. As further discussed in the section *Future Challenges*, there are a lot of remaining challenges to solve on this question.

**Fig.2.**
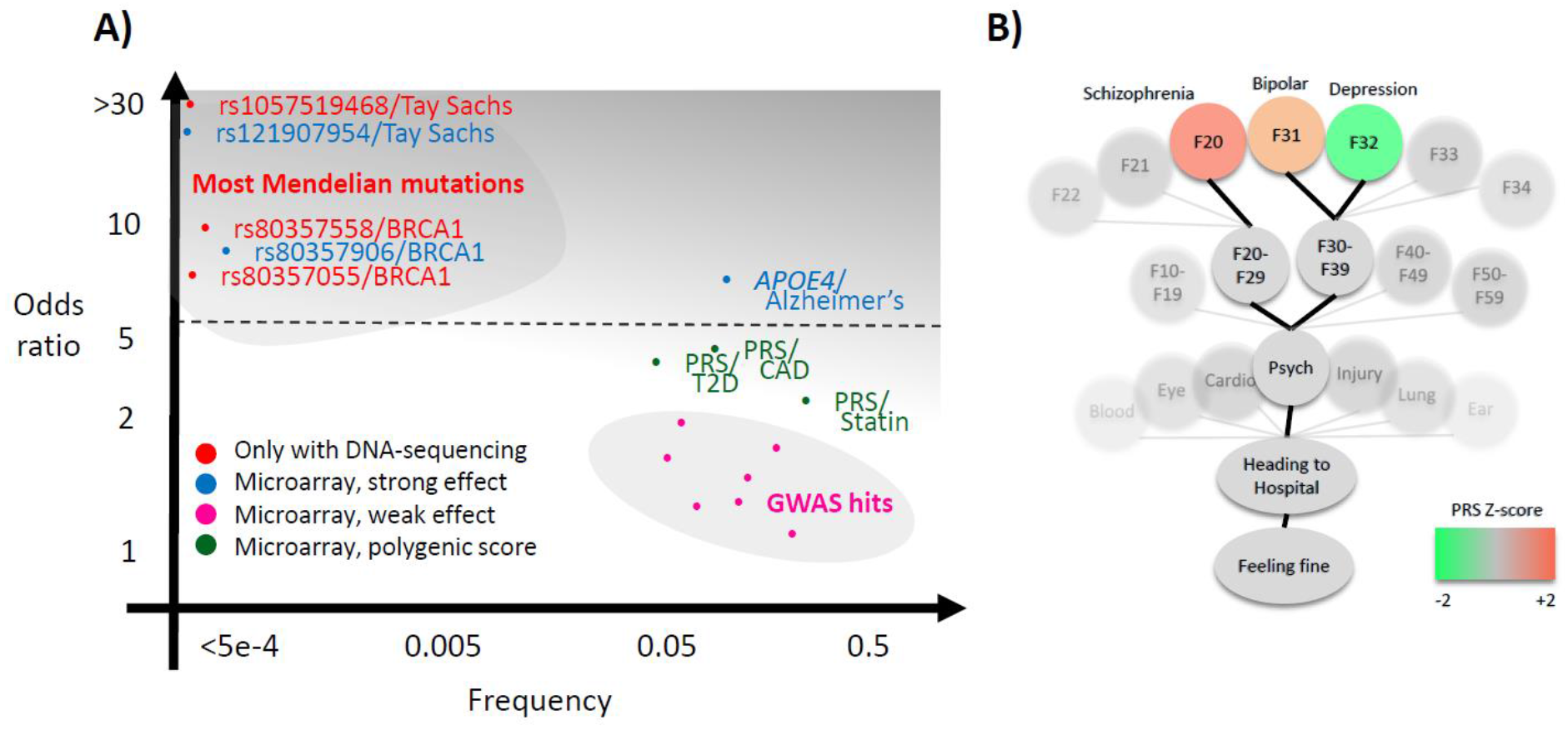
Theoretical background of the analysis pipeline. **A) C**linical genetics currently concern high-effect DNA-variants that often can only be sequenced (*red*). Additionally, high-effect variants such as APOE4 and a small subset of BRCA1 and BRCA2 pathogenic variants are possible to measure using microarray (*blue*, includes several other variants not shown in plot, e.g. Parkinson’s variants). There may be an untapped potential for valuable clinical information in polygenic risk scores (PRS) for common disease (*green*), for example for type 2 diabetes (T2D), coronary artery disease (CAD) or Statin response [Natarajan et al 2017, Wünnemann et al 2019, Khera et al 2018]. It is a primary aim of the impute.me project to make this potential available more broadly, balancing the practice of relying on individual GWAS SNPs and/or reporting of single SNP genotypes (*pink*). **B)** The secondary aim is to provide genetic scores in a relevant context, exemplified in the precision medicine module showing the so-called health-context tree. This tree consists of all entries from the international classification of disease (ICD-10), linked to all genetic studies. It allows browsing of polygenic risk scores in a relevant context. In the example shown, the tree is open on the psychiatry chapter, showing polygenic risk scores for schizophrenia (F20), unipolar depression (F32) and bipolar depression (F31). While these scores have little predictive relevance for a healthy individual, they may be useful in the context of psychiatric evaluation, particularly in the case of more extreme scores.

## 3 Validation and testing

To evaluate pipelines on individuals with known disease outcomes, we investigated 242 samples from the CommonMind data set. The CommonMind data set includes patients with schizophrenia, bipolar disorder and controls, from European ancestry and from African ancestry. For each disorder and each ancestry group the full impute.me pipelines were applied, including imputation and PRS-calculation. Additionally, SNP-sets corresponding to each of three major DTC companies were extracted and re-calculated. This was done to test the hypothesis that PRS calculation in mixed SNP-sets poses particular challenges with regards to missing SNPs. Such sets of genotyped SNPs that are different in each sample, is an unavoidable consequence of working with online data uploads.

We found that disease prediction strength, measured as variability explained, corresponded well to theoretical expectations of known SNP heritability [Li et al 2017, Wünnemann et al 2019]. Secondly, we found that using all-SNP scores resulted in better prediction than top-SNP scores, which was as expected [Vilhjálmsson et al 2015]. Thirdly, we found that prediction was more accurate in individuals of European ancestry compared with individuals of African ancestry, which is concordant with the PRS being developed from European Ancestry GWAS [Hou et al 2016, PGC 2014]. These observations match well with findings from studies of PRS in much larger data sets. Estimates of variability explained are probably more precise when derived from larger and more balanced studies, but the intention here is to provide a specific test of impute-me pipelines and address DTC-data related questions.

Of importance to this, we found that PRS prediction in mixed samples of non-imputed data causes severe problems. When training PRS algorithms, a SNP set is pre-specified. The pipelines evaluated here were trained with HapMap3 as SNP-set. Similar choices are made in other published PRS. However, such SNP-sets may not match with the SNPs available in downloadable raw data from DTC vendors. We therefore tested what prediction strength would be possible when using raw data directly from DTC vendors, both in a uniform setting (e.g. “all individuals use 23andme v4 data”) and in a mixed setting (e.g. “individuals have data from different vendors”). We found that in the uniform setting roughly half the predictive strength remained when using genotype data that is not imputed to match the HapMap3 SNP-sets (figure 3, row 2 and 4). In the mixed setting, virtually no predictive strength remained (figure 3, row 3 and 6). The mixed setting is the reality that is faced, both for third-party analytical services but also for DTC vendors with different chip versions. Imputation is therefore likely to be an essential requirement in such scenarios.

**Fig 3.**
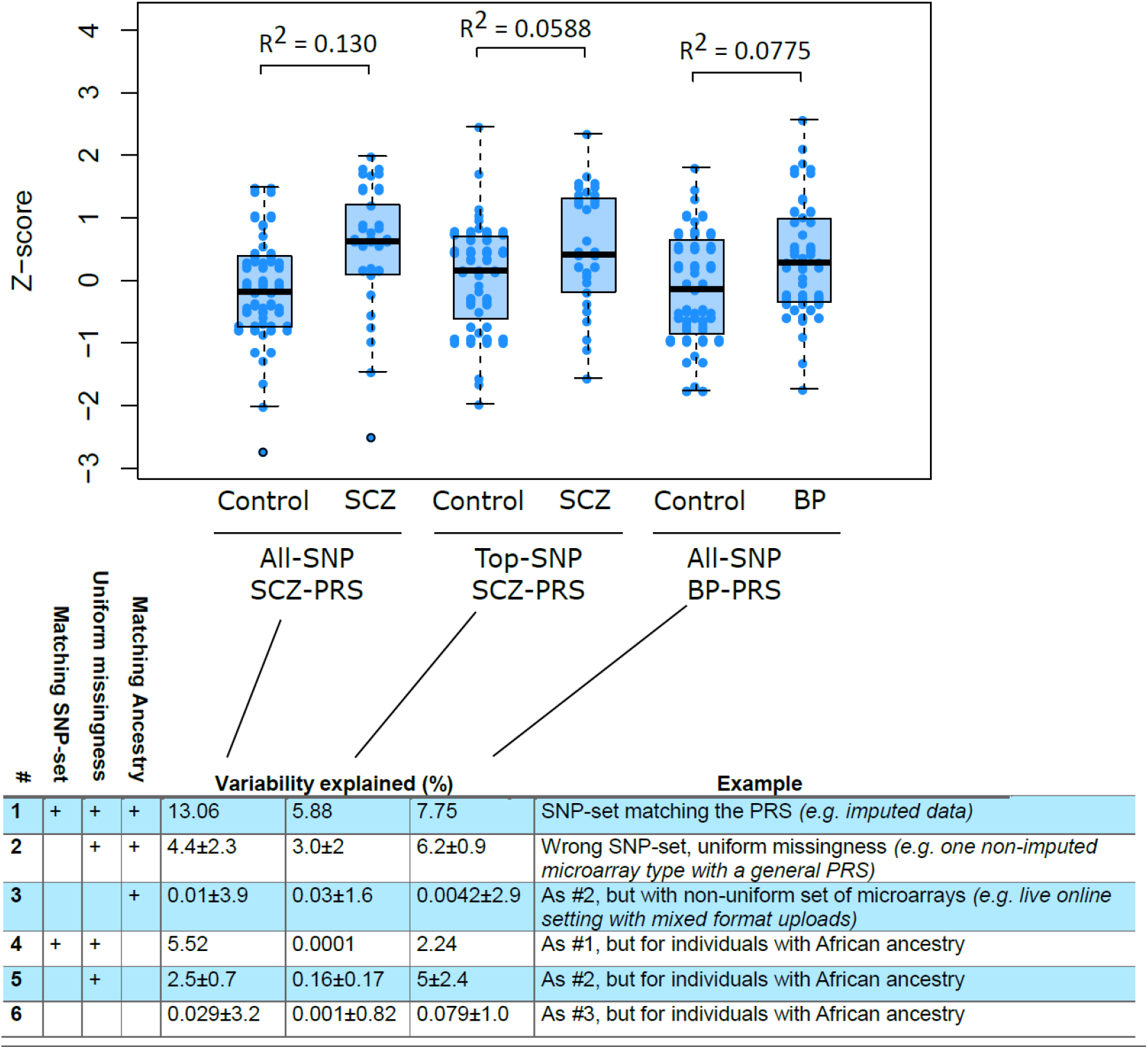
Pipeline evaluation using publicly available genotyped cohorts. Three scores were calculated in individuals of European ancestry and relevant diagnoses (n_control_=39, n_SCZ_=25 and n_BP_ = 39): a schizophrenia (SCZ) all-SNP score (n_SNP_=558406), a SCZ top-SNP score (n_SNP_=93) and a bipolar (BP) all-SNP score (n_SNP_= 554977). The BP top-SNP score only used 5 genome-wide significant SNPs and was not tested. The proportion of variance explained (Nagelkerke R^2^) is shown above each case-control pair. **B)** Testing different conditions of ancestry and input SNP-sets. Row #1 corresponds to the variability explained after processing through the full impute.me pipeline, i.e the same calculation shown in the plot. Row #2 shows the prediction level when the PRS algorithm uses input samples from only one type of direct-to-consumer (DTC) vendor, but the algorithm has not been trained specifically for that SNP-set. Values are given as mean±SD of three analyses in which SNP-sets were all from either 23andme (v4), ancestry-com, or myheritage, respectively. Row #3 shows the prediction when each sample uses different SNP-sets, i.e. the actual situation when dealing with user-uploaded DTC data online. Values are given as mean±SD over hundred random drawings of combinations of the 23andme (v4), ancestrycom, and myheritage sets, in proportions of 55%, 30%, 15%, respectively. These proportions correspond to what is observed in live users. Rows #4-6 shows the same as #1-3 but calculated for CommonMind individuals of African ancestry (n_control_=47, n_SCZ_=39 and n_BP_ = 6). The corresponding AUC values for this figure is 0.693, 0.614, 0.634 for row #1. For row #2: 0.55±0.12, 0.53±0.084, 0.62±0.012 and for row #3 0.58±0.047, 0.55±0.03, 0.57±0.047. Additionally, an extended version of the figure is available at www.impute.me/prsExplainer where additional metrics of prediction can be explored interactively.

To compare these findings with approaches that look at one SNP at the time, we extracted the SNPedia/Promethease SNPs that were indicated as associated with schizophrenia [Cariaso et al 2011]. All cases (n=25) and all controls (n=39) had at least one risk-variant from at least one of the 139 SNPs indicated schizophrenia associated. When focusing on SNPs that had the SNPedia/Promethease-defined “*magnitude*”-level (sic.) at > 1.5, we found that 80% of the SCZ cases (20 of 25) had at least one SNPedia/Promethease risk variant. Among the healthy controls 84% (33 of 39) had at least one such risk variant (p=0.9 for difference in proportions). In other words, it is not very predictive to know if you have a schizophrenia SNP. This illustrates the importance of considering more than one SNP at the time.

Finally, we compared pipeline reproducibility using two genome-data files, one obtained from MyHeritage and one from Ancestry.com, but sampled from the same person. After processing through the impute-me pipelines the correlation between PRS values over 1468 traits was r=0.933 between the two samples. Traits that showed discrepancy between the two data files typically were based on only few SNPs, of which one did not meet imputation quality thresholds for one of the data files.

## 4 Evaluation of Usage

As of June 2019, a total of 28 651 genomes had been uploaded to impute.me, and a total of 3.1 million analytical queries had been performed (figure 4A). The following additional observations about user behaviour may be of use to the genetics research community.

**Fig 4.**
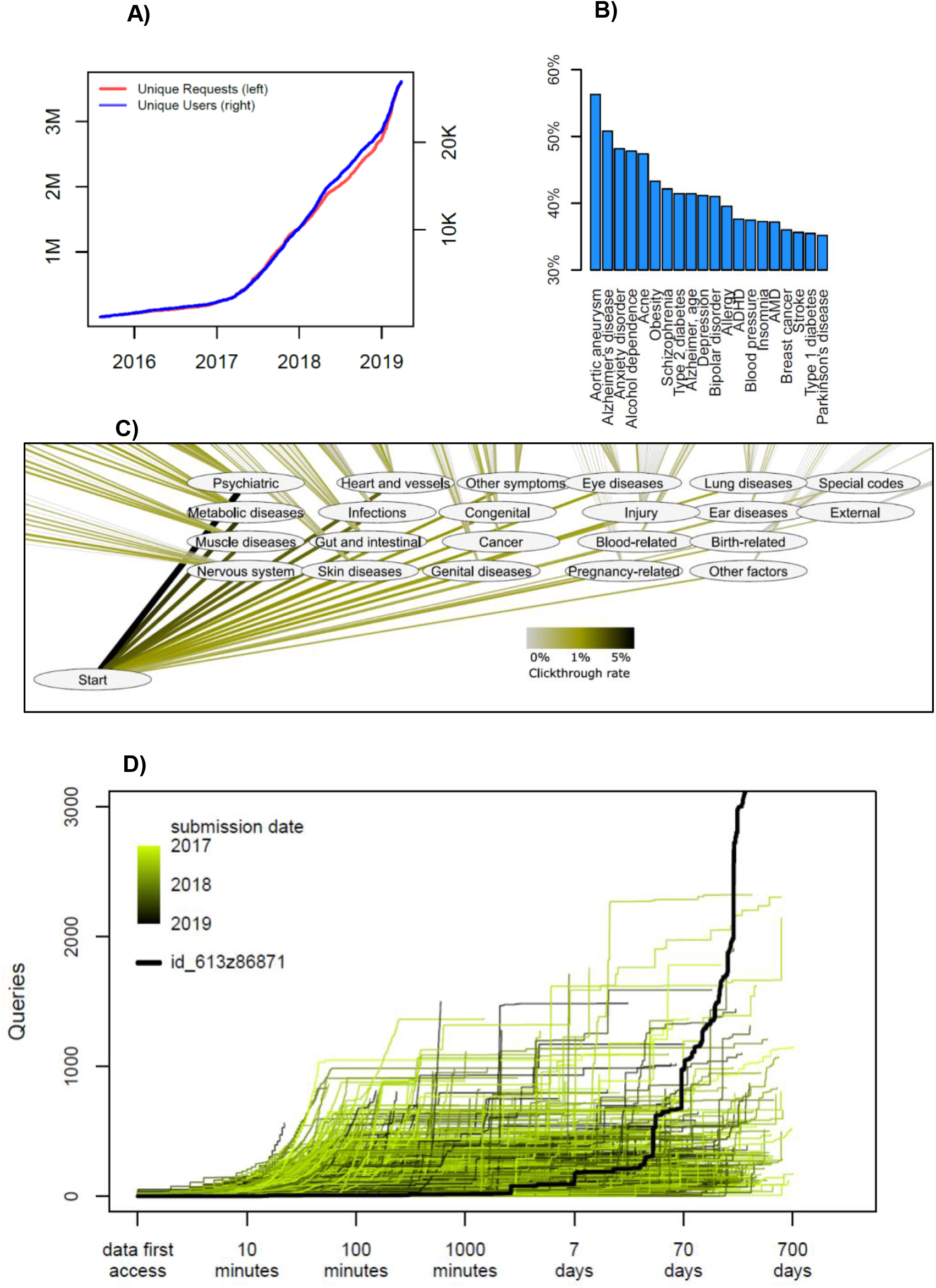
Detailed usage statistics. **A)** Overall count of unique users and unique analysis requests since august 2015. Each *request* corresponds to a specific analysis, e.g. the risk score for a disease, or a view in the ICD-10 based map in the precision medicine module. Each *user* corresponds to an uploaded genome with a unique md5sum. There is no check for twins, altered files or users with data from separate DTC companies. **B)** Distribution of user-interests in a trait in the *complex disease module*. In this module, each disease-entry is presented on an alphabetically sorted list, with aortic aneurysm being the default value. The percentage indicates how many of the users that scrolled down and selected this disease at least once (n_clicks_ = 871855). **C)** Distribution of interests in a trait in the *precision medicine module*. In this module, each disease-entry is presented in the layout of the ICD-10 classification system. The click through-rate reflects how many users pursued information in a given chapter or sub-chapter, as percentage of total amount of clicks (n_clicks_ = 114039). **D)** Analysis of how individuals use the interface over time. For each user, the number of queries is shown as a function of time after they first access their data. Since all data is automatically deleted after two years, no queries extend beyond 730 days. The colour code indicates the submission date. The highlighted black line indicates the publically available permanent test user with ID id_613z86871, which is omitted from all other analyses.

Common and well-known diseases are the most sought after. By overall click-count and comparing over several different modules, there is no doubt that users are most interested in common disease types; diseases of the brain, the heart and of the metabolism are more requested. Interface design may of course play important roles in such choices. For example, the choice to serve disease traits as alphabetically sorted lists is likely to artificially inflate interest in e.g. abdominal aneurysm (figure 4B). However, the larger interest in psychiatry, cardiovascular and metabolic disorders remain also in the precision medicine module which is not presented as an alphabetically sorted list (figure 4C). It is possible that greater scientific interest in PRS in these fields also drives some of these effects, but we cannot explain why other fields where PRS are actively discussed, such as cancer, are not attracting more attention

Likewise, it seems that common disease (“Complex disease module”) is more sought after than rare disease (“Rare disease module”); 95% of all users visit the first, whereas only 70% visit the second. Again, interface design and project goals probably play a big role in this – the landing page headers says *Beyond one SNP at the time* and the rare disease module is found in the navigation bar only below seven other module entries. But it may also illustrate a central communication challenge for the field: People are more interested in the genetics of common, complex disease with small effect sizes (figure 2A, lower right), but may interpret the results as if they were for rare disease with large effect sizes (figure 2A, upper left).

Finally, we have observed that usage of health genetic data surprisingly often is not just a test-and-forget-event. When plotting query-count as a function of time from first data-access, we find an expected pattern of intense browsing the hours and days after first data access (Figure 4D). However, many users re-visit their data even months and years after first data access, perhaps implying that results are considered and saved and then re-visited at a later time in a different context.

## 5 Future Directions

Generation of the PRS data presents one set of challenges but communicating them to people in such a way as to make it both comprehensible and useful presents another [Lipkus et al 1999, Naik et al 2012]. We believe that this is a crucial unmet need in current genetics research, because presenting PRS data in a way that is useful requires an understanding of people’s motivations for accessing them in the first place.

To date, studies of PRS have focused on providing people with PRS information in relation to specific conditions (e.g. cancer [Bancroft et al 2014, Bancroft et al 2015, Young et al 2017, Smit et al 2018]) for which participants have an indicated risk and exploring understanding and reactions. No studies have examined what motivates people to seek out and access their own PRS for common complex conditions, and little is known about how people understand or respond to the data they receive.

PRS information is inherently probabilistic in nature, which is well known to be difficult for people to understand [Hallowell et al 1998, Smerecnik et al 2009] and receiving information about genetic risk is not necessarily benign. When people receive genetic test results that they perceive to reflect high risk for a condition, this can have negative impact on outcomes like self-perception and affect, and in the case of receiving high risk test results for Alzheimer’s disease – can actually impact objective measures of cognitive performance [Wilhelm et al 2009, Dar-Nimrod et al 2013, Lineweaver et al 2014, Lebowitz et al 2017, Turnwald et al 2019]. Therefore, how information about genetic risk is communicated matters.

The literature suggests that when communicating risk, the most useful and effective strategy is to use absolute risks [Lipkus et al 1999, Naik et al 2012, Reyna et al 2009]. In the case of PRS with modest predictive power, however, this may simply result in re-stating the population prevalence of a disease for everyone [Janssens 2019]. It is therefore important that the predictive strength is also included in this communication; that is how much the genetic component potentially could alter the absolute risk. The genetic component corresponds to the SNP-heritability, and we are therefore exploring how to best include this information (e.g figure 1C). Currently we have registered the SNP-heritability for 294 of the reported traits, available as an experimental option called ‘plot heritability’. We believe that a main future direction is to experiment and expand on how to best communicate this to people.

It will therefore be useful to have a constant flow of people that are interested in interpreting their genetics and expose them to various modes of presentation. Some could involve statistically advanced concepts, like AUC and SNP-heritability, but others may take simpler approaches, such as the explanatory jar model pioneered for talking with families about genetics [Peay et al 2011, Austin et al 2019]. One may even imagine layered models of increasing complexity. This should be followed-up with questionnaires probing the level of understanding and general impact on users, something that is possible using the impute.me platform.

## 6 Conclusion

In summary, impute.me is a fully operational genetic analysis engine covering a very broad range of health-related traits, specifically focusing on optimizing possibilities from microarray-based DNA measurements. The challenges, their solutions, and the curation work behind them are highly relevant today in a setting of highly varying quality in interpretation of personal consumer genetics. They will also continue to be relevant in a future where we can expect both increases in the predictiveness of polygenic risk scores, as well as vast increases in the number of individuals buying consumer genetics. With a directed push towards responsible use of consumer genetics, this may even prove to be an overall clinical benefit.

For that reason, the impute.me code is available open-source open-source (see URLs) and the online implementation is free to use, supported by user-donations.

## 7 Methods

### 7.1 Pre-processing

Impute.me users upload raw genotype data from a DTC that is then parked in an imputation queue, which is simply a specific folder on our servers. Submitted jobs are then processed by a cron-job in order of free computing nodes becoming available. This cron-job consists of a series of calls to bash-based programs; first a shapeit call is made to phase the data correctly [Delaneau et al 2013], then an impute2 call is made with 1000 Genomes version 3 as reference [Howie et al 2009, 1000 genomes 2015]. Although the pipelines are not guaranteed to handle any format they receive, they currently operate with less than 1% processing failures, meaning uploads that cannot proceed through the full quality control and imputation pipelines. The failures are typically due to file formatting errors, missing chromosomes or any number of other odd data corruptions that real-world data exchange suffers from.

Several customizations have been made with the goal of minimizing memory foot print and thereby allowing running in a clustered fashion on a series of small cloud computers. This allows for relatively easy scaling of capacity; one simple setup (“hub-only”), where calculations are run on the same computer as the web-site interface. Another is a hub+node-setup, where a central hub server stores data and shows the web-site, while a scalable number of node-servers perform all computationally heavy calculations. After pre-processing is finished, two new files are created: a .gen file with probabilistic information from imputation calls and a *simple format* file with best guess genotypes, called at a 0.9 impute2 INFO threshold. All further calculations are based on these files.

### 7.2 Polygenic Risk Score Calculation

From the pre-processed data a modular set of trait predictor algorithms are applied. For many of the modules, the calculations are trivial. For example, this could be the reporting of presence and/or absence of a specific genotype, such as ACTN3 and ACE-gene SNPs known to be (weakly) associated with athletic performance. These are included mostly because users expect them to be. For other, we rely heavily on PRS.

An important distinguishing factor between different PRS algorithms is how risk alleles are selected. A commonly used approach includes variants based on whether they surpass a given p-value threshold in the GWAS, retaining only LD-independent variants using LD-based clumping, often with a p-value threshold of genome-wide significance (p<5e-8). Herein we refer to this approach as the ‘top-SNP’ approach. The top-SNP approach has the advantage that it is simple to explain, is easy to obtain for many GWAS, and has a light computational burden, e.g. [Buniello et al 2019, Watanabe et al 2019]. However, research has repeatedly shown that the inclusion of variants that do not achieve genome-wide significance improves the variance explained by PRS, with PRS including all variants often explaining the most variance. PRS based on GWAS effect sizes that have undergone shrinkage to account for the LD between variants have been shown to explain more variance than PRS which account for LD via LD-based clumping [Vilhjálmsson et al 2015]. Herein we refer to this approach as the all-SNP approach. It is more computationally and practically intensive to implement at scale. Consequently, within Impute.me each trait or disease reported shows all-SNP based PRS calculations if such is available, and top-SNP based PRS calculations if not.

In the top-SNP calculation mode, the results are scaled such that the mean of a population is zero and the standard deviation (SD) is 1, according to the relevant 1000 Genomes super-population; either African, Ad Mixed American, East Asian, European or South Asian.

Population-score_snp_ = frequency_snp_ * 2 * beta_snp_

Zero-centered-score =Σ Beta_snp_ * Effect-allele-count_snp_ - Population-scoresnp

Z-score = Zero-centered-score / Standard-deviation_population_

Where beta (or log(odds ratio)) is the reported effect size for the SNP effect allele, frequencySNP is the allele frequency for the effect allele, and the Effect-allele-countSNP is the allele count from genotype data (0, 1, or 2).

In the all-SNP calculation, the scaling is similar, but done empirically, i.e. based on previous Impute.me users of matching ethnicity. This mode of scaling is also available as an optional functionality in the top-SNP calculations, and generally seems to match well with the default 1000 genomes super population scaling.

The all-SNP scores were derived using weightings from the LDpred algorithm [Vilhjálmsson et al 2015]. This algorithm adjusts the effect of each SNP allele for those of other SNP alleles in linkage disequilibrium (LD) with it, and also takes into account the likelihood of a given allele to have a true effect according to a user-defined parameter, which here was taken as *wt1* i.e. the full set of SNPs. The algorithm was directed to use hapmap3 SNPs that had a minor allele frequency >0.05, Hardy-Weinberg equilibrium P>1e-05 and genotype-yield >0.95, consistent with our expectation that these would be the best imputed SNPs after full pipeline processing.

### 7.3 Pipeline Testing

The CommonMind genotypes measured with the microarray of the type H1M were downloaded along with phenotypic information. Each sample was processed through the impute.me pipelines, using the batch-upload functionality. Reported ethnicity was compared with pipeline (genotype) assigned ethnicity and found to be concordant.

After pipeline completion, we extracted three PRS for each sample, corresponding to SCZ all-SNP, SCZ top-SNP and BP all-SNP. In the github-repository for impute.me, these three corresponds to the scores labelled *SCZ_2014_PGC_EXCL_DK.EurUnrel.hapmap3.all.ldpred.effects*, *schizophrenia_25056061*, *BIP_2016_PGC.All.hapmap3.all.ldpred.effects* trait-IDs [PGC 2014, Hou et al 2016]. These extracted scores formed the basis of the row #1 and #4 calculations in figure 3. The remaining rows were created by subsetting the best guess imputed genotypes into new sets of users, corresponding to each of three major DTC vendors and then re-running the scoring algorithms either with uniform data or mixed data. Uniform data is here defined as all samples 195 samples having the same set of SNPs available, corresponding to one of three DTC vendors in each run. Mixed data is defined as samples having different sets of SNPs available, a set corresponding to actual distributions of customers from different DTC vendors, with distributions re-drawn 100 times. We estimated the predictive ability of the PRS using Nagelkerke’s *R*^2^ and AUC.

### 7.4 Usage evaluation

A log data freeze was performed 2019-06-08, by making a copy of all usage log files and then removing the uniqueID of each user. This was done to prevent it from being linked with the genetic data of that user. The exception was the publicly available permanent test user with ID id_613z86871, which was lifted out before analysis and is not included in other summary statistics. Generally, a user corresponds to an uploaded genome with a unique md5sum. Click-through rates were calculated as fraction of users that performed any query in the module in question, e.g. the precision medicine module was only launched in September 2018 and therefore only counts clicks from people who have used it. Plots were generated using base-R version 3.4.2 and cytoscape version 3.71.

## 10 Author Contributions

LF coded the code. All authors contributed to interpretation, drafting the work, critical revision for important intellectual content, as well as final approval of the manuscript.

## 12 Acknowledgments

In addition to the crucial resources cited in the text, we wish to thank the CommonMind Consortium for availability of testing data. CommonMind is supported by funding from Takeda Pharmaceuticals Company Limited, F. Hoffman-La Roche Ltd and NIH grants R01MH085542, R01MH093725, P50MH066392, P50MH080405, R01MH097276, RO1-MH-075916, P50M096891, P50MH084053S1, R37MH057881, AG02219, AG05138, MH06692, R01MH110921, R01MH109677, R01MH109897, U01MH103392, and contract HHSN271201300031C through IRP NIMH.

## 13 Manuscript contributions to the field (200 words required)

The manuscript describes an approach for obtaining polygenic risk scores (PRS) for any individual using direct-to-consumer genetics. PRS are considered state-of-the-art for genetic prediction in common, complex diseases, and the availability of such methods therefore have the potential expand the usage of genetics beyond rare mendelian disorders. At the same time, the usage and interpretation of direct-to-consumer genetics is by many considered a controversial issue that needs more regulatory oversight. By setting this type of information free in the form of open-source code and web-applications, this manuscript provides input to a debate on what constitutes beneficial use of genetics in common, common complex disease.

## 14 Data Availability Statement

Publicly available datasets were analyzed in this study. This data can be found here: CommonMind data DOI: 10.7303/syn2759792.

## 15 URLs

Code repository: https://github.com/lassefolkersen/impute-me

Web ressource: https://www.impute.me/

